# Framework for combined functional and computational assessment of variant pathogenicity in the sodium leak channel NALCN

**DOI:** 10.1101/2025.09.15.676065

**Authors:** C. Weidling, H. Harms, A. Ameen, A. Panfilova, T. Gloystein, A. Stein, H.C. Chua, S.A. Pless

## Abstract

The sodium leak channel nonselective (NALCN) is the pore-forming channel component of the NALCN channelosome. This protein complex generates a depolarizing sodium leak in various tissues and contributes to respiratory and circadian rhythms, locomotion, and sleep. *De novo* and inherited variants of *NALCN* can lead to severe developmental syndromes called ‘contractures of the limbs and face, hypotonia, and developmental delay’ (CLIFAHDD) and ‘infantile hypotonia with psychomotor retardation and characteristic facies 1’ (IHPRF1), respectively. Although variants of uncertain significance (VUS) or presumed pathogenic variants have been studied in heterologous expression systems before, there is no generally accepted framework on how to assess or predict variant pathogenicity. We set out use the functional and computational characterization of 19 VUS detected in CLIFAHDD and IHPRF1 patients to establish a robust analysis to classify suspected disease-causing variants. Specifically, we employ a combination of multiple parameters derived from two-electrode voltage-clamp electrophysiology recordings and predicted protein stability and conservation scores. We show that this approach is capable of distinguishing benign common variants from both gain- and loss-of-function (GoF/LoF) variants. Additionally, our work provides mechanistic insight into the molecular mechanism underlying specific variants and provides insight into the unusual propensity of *NALCN* missense variants to result in GoF phenotypes. We anticipate that this experimental and computational framework will aid assessment of variant pathogenicity of NALCN and other components of the channelosome in the future.

## Background

The sodium leak channel nonselective (NALCN) is a widely expressed membrane protein that belongs to the superfamily of voltage-gated sodium and calcium channels (Nav/Cav) (Lee et al. 1999; Lu et al. 2007). Unlike Navs and Cavs, NALCN is encoded by a single gene and its constitutive activity requires assembly with the membrane protein FAM155A and two large intracellular proteins, UNC79 and UNC80. Together, these four proteins form the NALCN channelosome (Chua et al. 2020; Bouasse et al. 2019; Kschonsak et al. 2022; Kang and Chen 2022; Zhou et al. 2022) (Fig 1A). The resulting protein complex gives rise to a sodium leak current that is highly sensitive towards direct pore block by extracellular divalent cations (Chua et al. 2020). The NALCN-mediated current depolarizes the resting membrane potential and plays an important role in many central physiological processes, including respiratory and circadian rhythms, locomotion, and sleep (Monteil et al. 2023).

**Figure 1:**
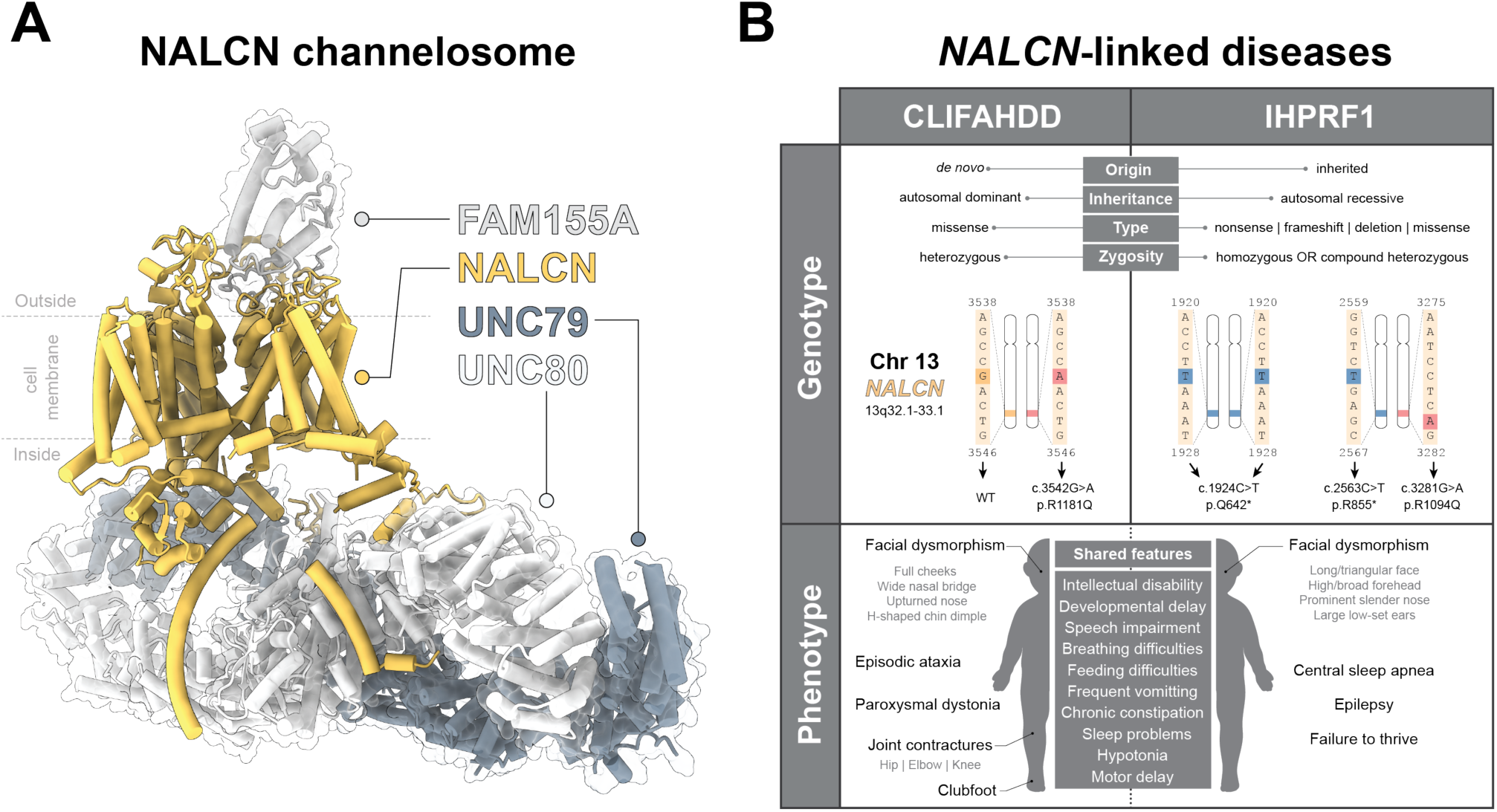
Overview of the NALCN channelosome and NALCN-linked diseases. (A) Structure of the NALCN channelosome, consisting of the pore-forming NALCN subunit along with auxiliary proteins UNC79, UNC80 and FAM155A (PDB 7SX3). (B) Genotypic origins and phenotypes of CLIFAHDD and IHPRF1.

Genetic variation associated with components of the NALCN channelosome can lead to rare but severe disease phenotypes. For example, heterozygous *de novo* variants of *NALCN* can lead to a severe developmental syndrome called ‘contractures of the limbs and face, hypotonia, and developmental delay’ (CLIFAHDD, OMIM #616226), an autosomal dominant syndrome caused by missense variants (Monteil et al. 2023; Chong et al. 2015) (Fig 1B). By contrast, inherited homozygous or compound-heterozygous variants of *NALCN* can cause ‘infantile hypotonia with psychomotor retardation and characteristic facies 1’ (IHPRF1 (OMIM #615419), an autosomal recessive syndrome caused by missense or nonsense variants, as well as frameshifts or deletions (Monteil et al. 2023; Köroğlu et al. 2013; Al-Sayed et al. 2013) (Fig 1B).

Many of the phenotypic manifestations of these ultra-rare syndromes, such as developmental delay, hypotonia, and feeding and sleeping difficulties are shared between CLIFAHDD and IHPRF1. Distal arthrogryposis, episodic ataxia, and paroxysmal dystonia are only observed in CLIFAHDD patients, who also exhibit more severe respiratory symptoms at birth. Failure to thrive, central sleep apnea, and epilepsy are more common among IHPRF1 patients (Fig 1) (Parra-Díaz et al. 2025). Nonetheless, efforts to create reliable genotype-phenotype relations are crucial for diagnostic and potential therapeutic purposes, but have proven difficult to establish (Parra-Díaz et al. 2025; Monteil et al. 2023). This is partly due to the low number of patients (<150 worldwide) and the lack of a robust and widely accepted methodological approach to test and classify putatively disease-causing variants in *in vitro* or *in vivo* models. Given the increasing number of reported NALCN-linked VUS (>400 in ClinVar alone), and the associated societal and personal burdens associated with such classifications, we set out to implement a functional *in vitro* assay for NALCN variants following the well-established guidelines by the American College of Medical Genetics and Genomics (ACMG, (Brnich et al. 2019)) to assess variant pathogenicity. The guidelines recommend a careful calibration of the functional test applied to putative disease-causing variants against common and likely benign protein variants, an approach that is validated for related Nav and Kv channels (Ma et al. 2024; Vanoye et al. 2022; Thompson et al. 2023).

We showed that assessing a combination of functional parameters in an electrophysiological *in vitro* assay can reliably discriminate between variants that - based on their common occurrence in the population – are likely benign, and a range of established or previously untested CLIFAHDD and IHPRF1 variants. In total, we tested four common variants, seven suspected CLIFAHDD variants, and 12 clinically characterized IHPRF1 variants. We demonstrated the power and versatility of this approach by identifying the underlying molecular mechanism of action of some variants, as well as re-classifying a suspected disease-causing variant as likely benign. Lastly, we used complementary computational approaches based on population sequencing data, structure-based predictions of mutation-induced changes in protein stability, and sequence tolerance estimated by large language models. These methods can cluster variants in agreement with our functional data and, in combination with it, can increase the confidence with which pathogenic variants can be identified and characterized.

## Methods

### Plasmid generation and RNA synthesis

Custom-designed oligonucleotides (Eurofins Genomics) were used for PCR-based site-directed mutagenesis using either the Q5 system (New England Biolabs) or PfuUltra II Fusion Polymerase (Agilent Technologies). Mutations were introduced into a modified pcDNA3.1(+) vector with codon-optimized NALCN/ NALCN-eGFP-2xFLAG tag (Chua et al., 2020). PCR products were analyzed on 1% agarose (Fisher Scientific, dissolved in TAE buffer) gels stained with SybrSafe (Fisher Scientific), at 100 V for 30-40 min. PCR products were digested with DpnI (New England Biolabs) to remove the template and transformed by heat shock into chemically competent *E. coli* XL10-GOLD cells, grown on LB agar, supplemented with 1 μg/μl Ampicillin. Plasmids were purified from liquid cultures using the Presto mini plasmid kit (Geneaid). Plasmids were sequenced (Eurofins Genomics/ Macrogen) throughout the full-length of the coding gene to confirm the mutation. For oocyte expression, plasmids were linearized with XbaI (New England Biolabs) at 37°C overnight and purified either using the GenepHlow Gel / PCR kit (Geneaid) or by EtOH precipitation: 1:10 volumes of 3 M NaOAc were added to stop the reaction, followed by 2.5 volumes of ice-cold 96% EtOH, after which the samples were incubated at -80°C for > 30 minutes. After centrifugation for 30 min at 16,000 x g at 4°C, the DNA was washed with cold 70% EtOH and then dried and re-suspended in nuclease-free water. DNA was transcribed into RNA using the T7 mMessage mMachine kit (Thermofisher), following the instructions of the manufacturer. RNA was purified by LiCl precipitation: 3 volumes of LiCl were added and the reaction was incubated at -20°C for at least 30 min, followed by washing twice with 10 volumes of ice-cold 70% EtOH. The RNA was air-dried and reconstituted in RNase-free water and concentrations were measured using a NanoDrop spectrophotometer (ThermoFisher Scientific).

### Xenopus laevis oocyte culture and injection

Xenopus laevis oocytes were prepared as described before (Chua et al. 2020) and healthy stage V-VI oocytes were injected with 1 µg/µl of pre-mixed RNA (18 - 46 nl of 1:1:1:1 plasmid ratio of NALCN/ respective mutant, UNC80, UNC79 and FAM155A). Cells were incubated in OR3 (50 % (v/v) Leibovitz’s L-15 medium (Gibco), 1 mM glutamine, 250 μg/mL gentamycin, 15 mM HEPES; pH 7.6 with NaOH) or ND96 supplemented with antibiotics (96 mM NaCl, 2 mM KCl, 1 mM MgCl2, 5 mM HEPES, 1.8 mM CaCl2, 2.5 mM sodium pyruvate, 0.5 mM theophylline, 0.05 mg / mL gentamycin, 0.05 mg / mL tetracycline; pH 7.4 with NaOH / HCl) at 18°C with agitation for 3-5 days post-injection.

### TEVC recordings and data analysis

Microelectrodes were pulled to resistances of 0.2-1.2 MΩ from borosilicate glass (Harvard Apparatus) using a Model P-1000 Flaming/Brown Micropipette puller (Sutter Instruments) and filled with 3 M KCl. Oocyte recordings were acquired using a Warner OC-725C oocyte clamp amplifier (Warner Instruments) and digitized using an AxonTM Digidata® 1550 system (Molecular devices), filtered (Hum Bug, Quest Scientific), and acquired / analyzed using the pCLAMP 10/11 software (Molecular Devices).

Cells were perfused with ND96 (96 mM NaCl, 2 mM KCl, 1 mM MgCl_2_, 1.8 mM CaCl_2_, 5 mM HEPES; pH 7.4 with NaOH / HCl) or Ca^2+^/Mg^2+^-free ND96 (96 mM NaCl, 2 mM KCl, 1.8 mM BaCl_2_, 5 mM HEPES; pH 7.4 with NaOH / HCl) in custom-made oocyte chambers during recordings and subjected to 1 s sweeps of +80 mV to -100 mV in decrements of 20 mV, from a holding potential of 0 mV. Data was analyzed from a minimum of six cells from two independent experiments for each tested construct and WT-injected cells were recorded from each batch for normalization. Raw data was subjected to 5× data reduction for visualization of the traces.

To plot the fold change in out- and inward current of the mutants, data were normalized to the WT average maximum current of each recording day. Significance of the fold change was analyzed by two-tailed, unpaired student t-test of each mutant to WT of the same recording days (absolute values) or one-way ANOVA followed by Dunnett’s multiple comparison. All recording data was analyzed using GraphPad Prism 9 and visualized in Adobe Illustrator 2020. All error bars are shown as mean ± standard deviation (SD).

### ΔΔG and sequence tolerance value prediction

Predicted thermodynamic stability values (ΔΔG) were calculated using the “cart_prot” protocol (Tiemann et al. 2023). We used the pipeline available at https://github.com/KULL-Centre/PRISM/tree/main/software/rosetta_ddG_pipeline (release tag v0.2.6, GitHub sha1 a6523b7438ee341b312cf77bda6378d5c3a74b0e) in membrane protein, saturation mutagenesis modes, with Rosetta version v2021.31-dev61729 (GitHub sha1 c7009b3115c22daa9efe2805d9d1ebba08426a54). All non-residue atoms were removed from the input structure (PDB ID 7SX4) at the cleaning step, and all protein chains were kept. The command used is: python3 ∼/PRISM/software/rosetta_ddG_pipeline/run_pipeline.py-s7sx4.pdb-o {output_path} -i fullrun -mm all --chainid A --run_struc ABCDE --overwrite_path True --slurm_partition {hpc_partition} --is_membrane True --mp_align_ref 7sx4_A

Predicted sequence tolerance values were provided by colleagues and obtained by them from ESM-1b as described (Cagiada et al. 2025). Data for NALCN and other human proteins can be retrieved at Electronic Research Data Archive at University of Copenhagen (ERDA, https://sid.erda.dk/cgi-sid/ls.py?share_id=DUWFpyjZp0) or using a Colab notebook (https://colab.research.google.com/github/KULL-Centre/_2024_cagiada-jonsson-func/blob/main/FunC_ESMs.ipynb). UniProt ID used to get NALCN scores is Q8IZF0.

Both stability prediction and sequence tolerance scores are output in-house file format containing protein sequence metadata, allowing for straightforward alignment of the predictions from different sources. We used an in-house parser (https://github.com/KULL-Centre/PrismData) to align ΔΔG values to the UniProt sequence, as the crystal structure used as an input for the ΔΔG calculations has several unresolved regions, and ESM1b is a sequence-based method that predicts scores for the entire UniProt protein sequence.

## Results

### CLIFAHDD-associated NALCN variants lead to gain-of-function via different mechanisms

We and others have previously characterized several CLIFAHDD-associated *NALCN* variants in heterologous expression systems (Kschonsak et al. 2020; Bouasse et al. 2019; Hadouiri et al. 2025) and *C. elegans* (Bend et al. 2016; Aoyagi et al. 2015) and found that they generally displayed a gain-of-function (GoF) phenotype, meaning that they caused the channel to be overactive relative to wild-type (WT) function. Since then, multiple new suspected CLIFAHDD variants have been reported in the literature. Similar to previously described variants, these novel variants also cluster around the pore domain (PD) and intracellular linkers of NALCN (Fig 2A), raising the question of whether they affect NALCN function in a similar way.

**Figure 2:**
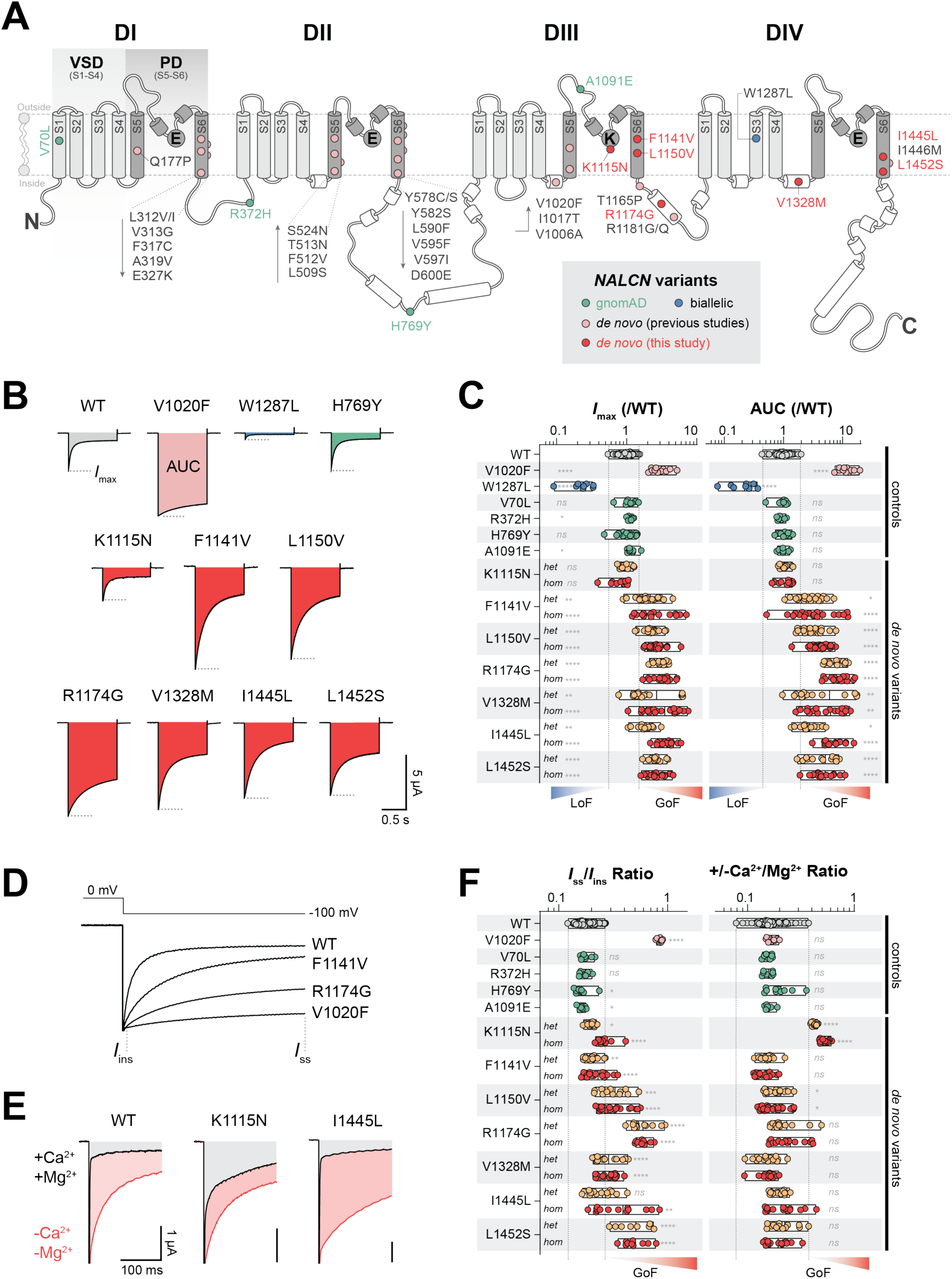
Functional characterization of CLIFAHDD variants consistently reveals GoF phenotypes. (A) A two-dimensional representation of NALCN structure highlighting the four homologous domains (DI-IV), each containing six transmembrane segments (S1-S6). The S1 to S4 segments of each domain make up the voltage-sensing domain (VSD), whereas the S5 and S6 segments, linked together by pore loops, form the pore domain (PD). The locations of common population variants found in the gnomAD population (green), the previously characterized only biallelic IHPRF1 missense variant (blue), as well as previously studied and novel de novo CLIFAHDD variants (pink and red, respectively) are indicated on the cartoon. (B) Representative current traces of NALCN WT and variants in response to a hyperpolarizing step (0 to -100 mV) in the absence of Ca^2+^ and Mg^2+^. Maximal currents (Imax) are indicated using grey dotted lines, whereas total charge movement is quantified as area under the curve (AUC), shown as colored fills. (C) Normalized maximal current amplitudes (Imax, left) and area under the curve (AUC) values (right) of NALCN variants (against WT mean of individual batches). (D) Overlay of current responses at -100 mV for WT and variants to highlight the differences in the instantaneous (Iins) and steady-state (Iss) components. (E) Representative current traces of NALCN WT and variants in the presence (+Ca^2+^+Mg^2+^) and absence (-Ca^2+^-Mg^2+^) of divalent cations. (F) Iss/Iins (left) and +/-Ca^2+^/Mg^2+^ (right) ratios for WT and variants. Data are shown as mean with min to max (floating bars). For controls (V1020F, W1287L, and population variants), unpaired t-test was used for statistical comparisons (against WT). For novel de novo variants, one-way analysis of variance (ANOVA), Dunnett’s test (against WT) was used. ns: non-significant, p>0.05; *p < 0.05; **p<0.01; ***p<0.001; ****p < 0.0001. het: heterozygous, hom: homozygous.

Here, we set out to characterize a set of seven of these previously untested *NALCN* variants by expressing them in *Xenopus laevis* oocytes together with the other components of the NALCN channelosome (UNC79, UNC80, and FAM155A) and testing variant function by two-electrode voltage-clamp (TEVC). Specifically, this included K1115N in the selectivity filter, F1141V and L1150V in S6 of DIII, R1174G in the DIII-DIV linker, V1328M just upstream of S5 in DIV, and I1445L and 1452S in S6 of DIV (myGENE2.org) (Bramswig et al. 2018; Vivero et al. 2017; Borovikov et al. 2019) (Fig. 2A). Initially, we used two parameters to characterize the basic function of NALCN in response to a voltage step from 0 mV to -100 mV: i) the maximum inward current (I_max_) and ii) the total charge passing through the channel during a 1-s hyperpolarization, measured as the area under the curve (AUC), each normalized to WT. To validate our assay, we first tested the previously characterized V1020F (GoF) and W1287L (LoF) variants (Bouasse et al. 2019; Kschonsak et al. 2020), which displayed robustly increased and decreased I_max_ and AUC, respectively (Fig. 2B, C). Additionally, we tested four common *NALCN* variants, selected based on their gnomAD allele frequencies (>0.001, given in parentheses for each variant): V70L in the N-terminal domain (0.00166), R372H in the DI-DII linker (0.00116), H769Y in the DII-DIII linker (0.0251), and A1091E in the pore loop of DIII (0.001099) (Fig. 2A). Consistent with the general expectation that common population variants are likely to be functionally neutral or benign, these variants displayed *I*_max_ and AUC values indistinguishable from WT (Fig. 2B, C, Table S1). These findings demonstrate that our assay has the dynamic range to confidently distinguish between various functional phenotypes, ranging from LoF, to WT-like, to GoF.

Testing the suspected CLIFAHDD variants, we found that six out of seven variants displayed varying degrees of increased I_max_ (2- to 3-fold on average) and AUC (3- to 8-fold on average), clearly indicating them as GoF variants (Fig. 2B, C). K1115N was a notable exception to this trend, showing both WT-like I_max_ and AUC. Because all currently known CLIFAHDD variants occur heterozygously in patients, we also tested the variants in a quasi-heterozygous condition. For this, we injected *Xenopus laevis* oocytes with a 1:1 mix of mRNA encoding for WT and variant NALCN (in addition to UNC79, UNC80, and FAM155A). Six out of seven variants displayed similarly increased I_max_ and AUC between their homozygous and heterozygous conditions (Fig. 2C). While observed in a rather simple model system, this dominant behavior may explain why a single variant allele is sufficient to cause a pathogenic phenotype in patients.

As heterologous expression of NALCN, and thus a detailed functional characterization, has only been achieved recently, we wanted to explore further how these variants affect other functional parameters beyond maximal current levels. To do this, we assessed two additional functional parameters: i) the ratio between steady-state and instantaneous current (*I*_ss_ / *I*_ins_ ratio) of the inward current at -100 mV (Fig. 2D) and ii) the ratio of I_max_ recorded in the presence and absence of extracellular Ca^2+^ and Mg^2+^ (+/-Ca^2+^/Mg^2+^ ratio) (Fig. 2E). We have previously shown that hyperpolarization elicits large inward currents that deactivate rapidly (Chua et al. 2020). Consistent with this phenotype, WT channel has an *I*_ss_ / *I*_ins_ ratio of around 0.2. This ratio was generally increased for CLIFAHDD-associated variants in both homozygous and heterozygous conditions, indicating impaired current deactivation, although this effect was comparatively modest for K1115N, F1141V, and V1328M. By contrast, the K1115N variant was the only variant to display a reduced block by extracellular Ca^2+^ and Mg^2+^, as indicated by a significantly increased +/-Ca^2+^/Mg^2+^ ratio (Fig. 2F, Table S1).

### IHPRF1 homozygous nonsense variants in NALCN render the channelosome nonfunctional

Our study had thus far focused only on CLIFAHDD variants, where all but one of the 31 previously tested variants, as well as the 7 from this study, generate a GoF phenotype (Kschonsak et al. 2020; Bouasse et al. 2019) (Fig. 2). Yet it remained unclear what molecular phenotypes *NALCN* variants suspected to give rise to IHPRF1 (typically inherited as homozygous (biallelic) or compound-heterozygous variants) would result in. We thus turned towards assessing the functional impact of a subset of 7 clinically described IHPRF1-causing variants, including homozygous deletions (V891Sfs*2, ΔV956-L963 (Bramswig et al. 2018) and ΔV1020-R1054 (Nguyen et al. 2023)), premature stop codons that are spread throughout NALCN (W107*, W179*, Q642*, R1008*, Q1186* and R1384* (Bramswig et al. 2018; Seven et al. 2002; Köroğlu et al. 2013)), as well as a single missense variant (G883V (Tehrani Fateh et al. 2023)) (Fig. 3A). We expressed these NALCN variants together with FAM155A, UNC79 and UNC80 in *Xenopus laevis* oocytes and found that with the exception of G883V, all variants showed significantly reduced *I*_max_ and AUC (Fig. 3B, C). As a consequence, *I*_ss_ / *I*_ins_ and +/-Ca^2+^/Mg^2+^ratios could not be assessed.

**Figure 3:**
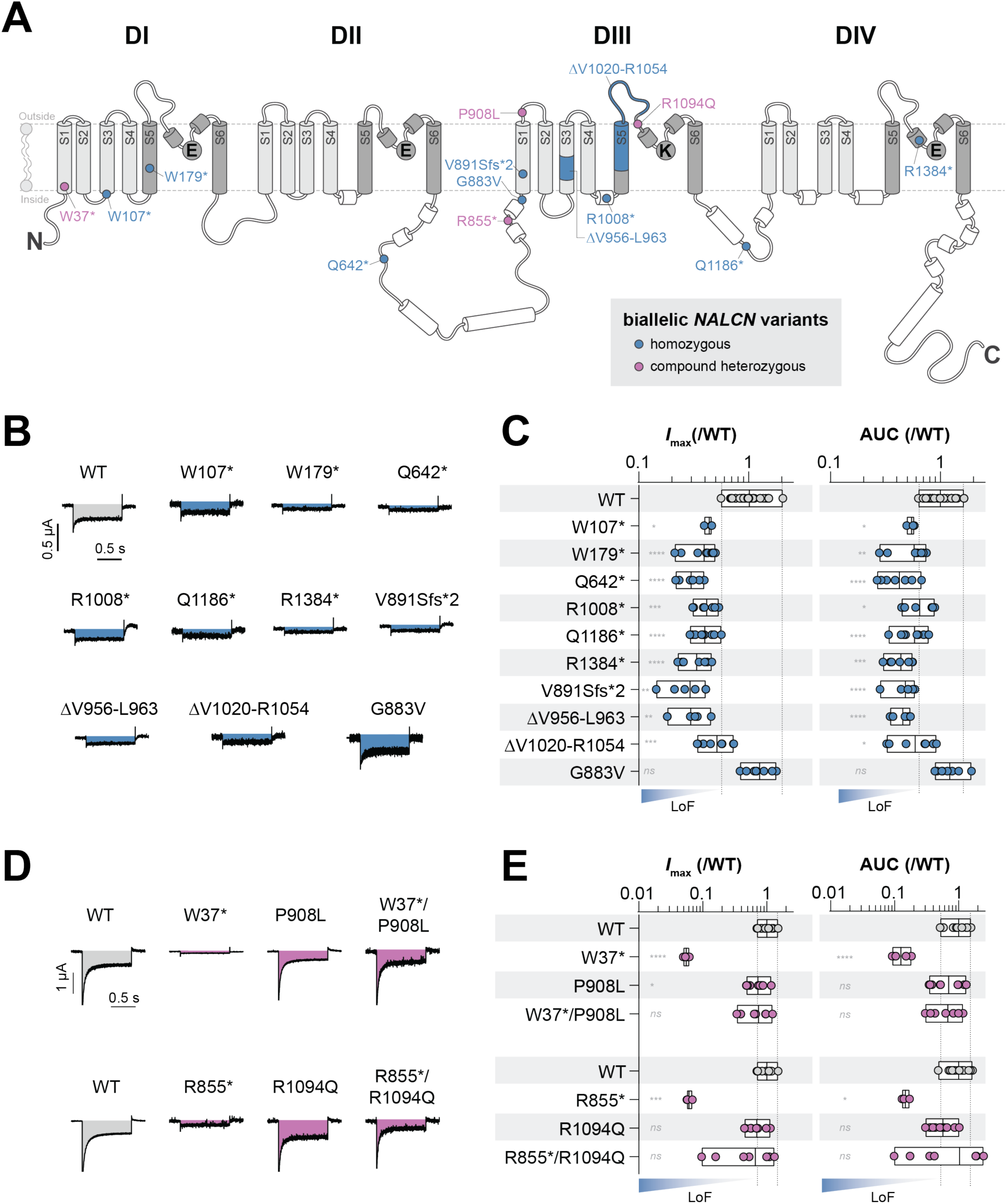
Functional characterization of IHRRF1 variants reveals predominantly LoF phenotypes. (A) A two-dimensional representation of NALCN structure, see Fig2A for details. The locations of homozygous (blue) and compound heterozygous (purple) IHPRF1 variants are indicated on the cartoon. (B) / (D) Representative current traces of NALCN WT and homozygous IHPRF1 variants (B) or compound heterozygous IHPRF1 variants (D) in response to a hyperpolarizing step (0 to -100 mV) in the absence of Ca^2+^ and Mg^2^+. (C) / (E) Normalized maximal current amplitudes (Imax) and area under the curve (AUC) values of NALCN variants shown in (B) / (D) (against WT mean of individual batches). Data are shown as mean with min to max (floating bars). For homozygous variants (B), unpaired t-test was used for statistical comparisons (against WT). For compound heterozygous variants (D), one-way analysis of variance (ANOVA), Dunnett’s test (against WT) was used. ns: non-significant, p>0.05; *p < 0.05; **p<0.01; ***p<0.001; ****p < 0.0001.

These findings demonstrate that homozygous internal deletions or large-scale truncations within the transmembrane or domain linker regions of NALCN are detrimental to the function of the NALCN channelosome. These results further mirror similar results obtained for IHPRF2-causing homozygous nonsense mutations in UNC80, which also display a pronounced LoF phenotype (Chua et al. 2020; Kschonsak et al. 2022). By contrast, we found no evidence that the homozygous G883V variant impacts the function of the NALCN channelosome. We therefore conclude that G883V is likely not pathogenic, while all variants with internal deletions or truncations (V891Sfs*2, ΔV956-L963, ΔV1020-R1054, W107*, W179*, Q642*, R1008*, Q1186*, and R1384*) result in LoF, consistent with being linked to IHPRF1.

### IHPRF1 compound heterozygous NALCN mutations result in mild LoF phenotypes

A recent study reported three IHPRF1 patients with relatively mild symptoms including developmental delay, hypotonia, and sleep apnea, and found the patients to carry two different combinations of compound heterozygous variants: W37* / P908L and R855* / R1094Q (Campbell et al. 2018) (Fig. 3A). We therefore set out to functionally investigate the effects of these variants alone and as combinations of nonsense and missense variants, resembling the respective compound heterozygous variant combination in patients.

As expected, we found that the nonsense variants W37* and R855* showed a marked LoF phenotype when expressed individually (Fig. 3B, C). This was evident from their dramatically reduced I_max_ and AUC values, which also prevented us from assessing I_ss_ / I_ins_ and +/-Ca^2+^/Mg^2+^ ratios. By contrast, currents from the missense variants P908L and R1094Q were WT-like in both I_max_ and AUC, and expression of W37* / P908L or R855* / R1094Q, resembling the variant combinations in patients, showed robust voltage-dependent currents with only minor reductions in I_max_ and AUC (Fig. 3D, E; Table S1).

It is also worth noting that three out of the four variants appear in the gnomAD database (allele count; R855*: 41; P908L: 13; R1094Q: 9), suggesting that they are less likely to be the monogenic cause of rare and severe diseases. Furthermore, patients carrying the W37* / P908L and R855* / R1094Q variants were also found to have additional variants that could be responsible for some, but likely not all symptoms reported in these patients (Campbell et al. 2018). As such, the pathogenicity of these NALCN variants remains uncertain.

### Computational analysis reveals variant clustering largely in line with functional results

Although firmly established, exclusively testing *NALCN* variants in simplified functional *in vitro* experiments, such as *Xenopus laevis* oocyte TEVC or patch-clamp electrophysiology assays, may still fail to correctly characterize a given variant as benign, LoF, or GoF, or lead to discrepancies with findings from other assays (Hadouiri et al. 2025). We therefore sought to increase the confidence in the predictive power of our functional findings by complementing them with an orthogonal computational approach. To this end, and to gain insights into possible protein-level changes of the tested *NALCN* variants, we integrated data from large-scale population sequencing (Karczewski et al. 2020), structure-based predictions of mutation-induced changes in stability (Park et al. 2016; Frenz et al. 2020), and sequence tolerance as estimated by a large language model (Rives et al. 2021). All three have previously been shown to correlate with experimental fitness of protein variants and distinguish disease-associated from neutral variants (Høie et al. 2022; Abildgaard et al. 2019; Cagiada et al. 2021; Larsen-Ledet et al. 2025).

In Fig. 4A we show the allele frequencies for NALCN mapped to its secondary structure, based on the AlphaFold2 model (Jumper et al. 2021) and in agreement with the available cryo-EM structures (Kang et al. 2020; Xie et al. 2020; Kschonsak et al. 2020). Regions of a gene or protein in which variants are depleted generally indicate functional relevance, whereas regions with greater variation are considered more tolerant to change and hence less functionally critical. While variants are distributed throughout the entirety of the NALCN protein, S5 and S6 of each domain, which form the ion-conducting pore, appear relatively depleted of variants (Fig. 4A). This aligns well with the observation that CLIFAHDD-associated variants typically occur around the pore of NALCN and generally result in severe GoF phenotypes (Kschonsak et al. 2020; Parra-Díaz et al. 2025; Bouasse et al. 2019) (Fig. 2). While GoF variants are typically difficult to predict with current computational tools, they have recently been reported to cluster structurally (Gerasimavicius and Marsh 2025), a notion that is supported by our results from NALCN CLIFAHDD variants.

**Figure 4:**
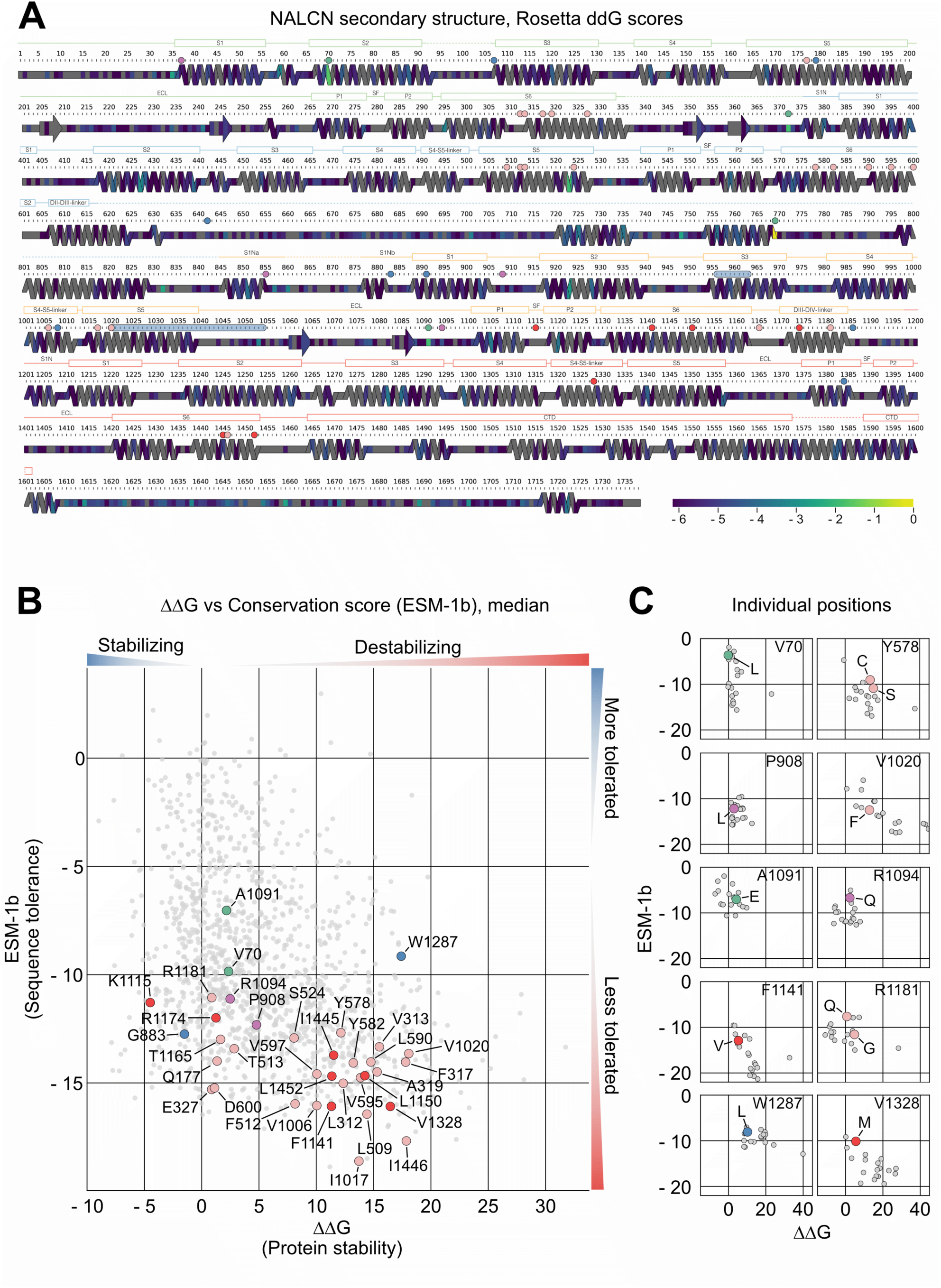
Protein stability predictions show variant clustering broadly in line with functional phenotypes. (A) Frequencies of genomic variants observed in NALCN in the gnomAD database (Karczewski et al. 2020). Color indicates the sum of variants observed at any given position; if there are different variants at the same position, their counts are added and divided by the total number of sequences available for the position. A log scale is applied to the frequencies to allow visualization of the broad range (gray: no variants are reported). Secondary structure is based on the AlphaFold2 model of NALCN and drawn with SSDraw (Chen and Porter 2023). (B) Scatter plot of the change in protein stability (ΔΔG in kcal/mol, x axis) and the estimated sequence tolerance (arbitrary units, y axis). By convention, positive ΔΔGs denote destabilizing variants, while negative ΔΔGs indicate stabilizing variants. Variants discussed in this manuscript are highlighted (green: common variants; pink: previously tested CLIFAHDD variants; red: CLIFAHDD variants tested in this study; blue: homozygous IHPRF1 variants; purple: compound heterozygous IHPRF1 variants). Variants close to 0 on both axes are expected to behave WT-like. Variants where mechanism of pathogenicity is based on loss of stability should be on the right side of the plot (ΔΔG > 3 kcal/mol), while in variants with ΔΔG near zero but low sequence tolerance scores, functional aspects such as interaction with other biomolecules are the likely cause of pathogenicity (Cagiada et al. 2021). (C) Scatter plots of the change in protein stability (ΔΔG in kcal/mol, x axis) and estimated sequence tolerance (arbitrary units, y axis) for individual positions highlighted in B. Colored data points denote amino acid exchanges that have been reported as common variants or patient variants (colors as in B), whereas grey data points represent all other possible exchanges for that position.

To evaluate the impact of variants on protein function, we used ESM-1b as a measure of estimated sequence tolerance and plotted it against ΔΔG as a measure of protein stability, as calculated by Rosetta (Tiemann et al. 2023). Loss of stability is commonly observed in pathogenic variants, with the loss of local or global protein stability leading to degradation of that protein variant. Sequence tolerance serves as a complementary measure, capturing similarities and discrepancies between orthologs of the protein family. Commonly observed variants (gnomAD allele frequencies >0.001) across the space of protein families are scored close to 0, while rare or unobserved changes score more negatively, indicating their likely detrimental nature. In a proteome-wide project (Cagiada et al. 2025), a threshold of - 6.5 was determined. It is, however, known that thresholds may vary by protein, just like different proteins are conserved to different levels, and as Fig. 4B indicates, pathogenic NALCN variants tend to have substantially lower scores than this general threshold. Values in Fig. 4B are displayed as the median of exchanges to every other amino acid for a given position, values of individual exchanges for a subset of positions are shown in Fig. 4C (values for all 35 highlighted positions can be found in Fig. S1). Of note, these calculations are limited to positions that are resolved in the currently available cryo-EM structures, which means that not all functionally tested variants could be plotted.

Values generally follow the expected distribution of more destabilizing exchanges (i.e. higher ΔΔG) displaying lower sequence tolerance (i.e. lower ESM-1b), but we further identify multiple notable trends: I) Positions of common variants show ΔΔG values close to 0, indicating limited impact in protein stability in line with their WT-like function, and relatively high ESM-1b values between - 5 and - 10, in agreement with their more frequent occurrence. For position V70 specifically, individual exchanges all score ΔΔG values close to 0, with V70L scoring among the best tolerated exchanges. However, both of these tendencies are less clear for A1091(E). II) Positions of CLIFAHDD-associated variants are more spread out compared to common variants, but mostly found at ΔΔG values above 10 and lower ESM-1b scores (< – 10), indicating stronger destabilization and lower sequence tolerance. An exception to this is R1181, including the most frequently found CLIFAHDD-associated variant R1181Q (>10 patients known worldwide). Despite an obvious GoF phenotype of R1181Q found by previous studies using different model systems (Kschonsak et al. 2020; Aoyagi et al. 2015), position R1181 displays a low ΔΔG value close to 0 and relatively high ESM-1b around – 11, positioning it closer to the commonly observed variants. Other recent works have highlighted challenges of current tools – including ESM – to identify GoF variants (Gerasimavicius and Marsh 2025). For many positions of CLIFAHDD variants, we find that individual exchanges display a rather wide spread of ΔΔG values. While this is also true for R1181, it is interesting to note that R1181Q specifically scores as the best tolerated exchange in its position. III) IHPRF1-associated compound-heterozygous missense variants display low ΔΔG (< 5) and relatively high ESM-1b around – 12, in line with their limited functional impact. Individual exchanges for both positions tend to score ΔΔG values close to 0, with both specific patient variants displaying ΔΔG values only slightly above 0. IV) Although based on only a single median data point, the position of the W1287L variant might indicate that IHRP1 partial LoF variants display comparatively high ΔΔG values combined with unusually high ESM-1b values. While this positioning is also observed for most individual exchanges in position W1287, identification and functional characterization of additional partial LoF variants is needed to confirm this notion.

In conclusion, and although not definitive, our computational analysis appears to be capable of broadly clustering variants. Importantly, the clustering is overall in agreement with the functionally observed phenotypes of the respective variants, thus serving as orthogonal confirmation.

## Discussion

In this study, we establish a combined ACMG-based functional and computational assay that is able to confidently discriminate between benign, LoF, and GoF variants of the NALCN channelosome. We then use this approach to characterize 19 suspected but previously not functionally tested CLIFAHDD and IHPRF1 variants. Adhering to this or similarly designed frameworks will establish the foundation for assisting clinicians and geneticists in more confidently assessing variant pathogenicity for NALCN in the future.

### Establishing a robust framework for NALCN variant classification

In the context of this study we achieve our goal of establishing a systematic and robust assay by means of three orthogonal and complementary approaches: First, guided by previously established ACMG guidelines (Brnich et al. 2019), we chose to select four *NALCN* variants that are amongst the most common in the general population (V70L, R372H, H769Y and A1091E) as controls. Indeed, these four variants display WT-like function in our *in vitro* functional assay. However, we were only able to validate our assay with two previously reported variants as LoF (W1287L) and GoF (V1020F), respectively, due to the novelty and rarity of pathogenic *NALCN* variants (Kschonsak et al. 2020; Bouasse et al. 2019). We recommend that the inclusion of multiple common variants, as well as at least one established GoF and LoF each be adopted as community standard to confirm the fidelity of the functional assay of choice. Generally, the scarcity of established variants as options for controls remains an issue, but this could potentially be offset, at least in part, through inclusion of variants such as R1181Q, for which multiple patient phenotypes are described and which has been characterized in multiple model systems (Monteil et al. 2023).

Second, we demonstrate the utility and necessity of assessing different metrics to test for altered function of the NALCN channelosome. In the past, we and others have primarily relied on comparisons of maximal inward or outward currents (*I*_max_), sometimes paired with biochemical assessments of (surface) expression levels (Kschonsak et al. 2020; Bouasse et al. 2019; Hadouiri et al. 2025). In other ion channels, this is often complemented by assessments of ligand sensitivity (EC_50_ in ligand-gated channels) or voltage dependence of activation/inactivation (V_1/2_ in voltage-gated channels), but this type of analysis is not reliable for a sodium leak channel with weak voltage sensitivity (Chua et al. 2020) and no known direct agonists. In this study, we therefore chose to not only compare *I*_max_ values, but also measures of total charge (AUC), current deactivation over time (*I*_ss_ / *I*_ins_), as well as sensitivity towards block by divalent cations (+/-Ca^2+^/Mg^2+^ ratio). Including these parameters expands the potential to identify defects in conduction, deactivation kinetics, and pore block and thus represents a potent means to identify and distinguish different molecular mechanisms underlying the pathogenic effect of the variants (see also below).

Finally, we capitalize on recent developments in computational methods to assess variant stability. Plotting ESM-1b conservation scores against ΔΔG protein stability predictions unveils reasonably defined variant clusters that broadly complement and support the results from the above functional assay. For example, the relatively common missense GoF variants typically have ESM-1b scores lower than -12, but across a relatively wide range of predicted protein stability values (-5 to 20). By contrast, functionally benign variants (such as common variants or compound-heterozygous missense variants found to be non-pathogenic), typically display ESM-1b scores between -5 and -12 across a narrow range of ΔΔG scores (around 0). Due to their rarity, missense LoF variants are more challenging to map, although the sole confirmed example (W1287L) appears to suggest localization around intermediate ESM-1B scores and high ΔΔG scores. Although we realize that implementing this exact assay as a complementary approach may not always be feasible, we recommend for the community to include, wherever possible, at least one orthogonal experimental or computational approach to validate findings from a given functional assay.

Although patient phenotypes will by nature always be more complex and variable than *in vitro* pathogenicity assessments (due to mosaicism, incomplete penetrance, epigenetics etc, see also (Hadouiri et al. 2025)), we propose that approaches such as our complementary functional and computational methods will serve as a benchmark for future studies of disease-causing variants of the components of the NALCN channelosome and possibly other proteins.

### Insight into disease-causing mechanisms

Our study supports the notion that homozygous nonsense variants, frameshifts, or deletions in NALCN result in complete LoF phenotypes. The molecular cause for their pathogenicity is relatively straightforward to rationalize based on the NALCN channelosome structure as any such alterations would either prevent assembly of the pore-forming NALCN protein or the association with the UNC79/UNC80 heterodimer. In fact, our previous work has shown that the NALCN C-terminus can be truncated after S6 of DIV (distal of position 1570 out of 1738) without major functional impairment, but that earlier truncations are detrimental (Chua et al. 2020). By contrast, UNC80 tolerates N-terminal deletions of around 700 out of its 3258 amino acids, while smaller C-terminal deletions would abrogate function, presumably because this would disrupt the interaction with UNC79 (Kschonsak et al. 2022).

All suspected CLIFAHDD variants tested here result in GoF phenotypes, both when expressed alone or in a quasi-heterozygous setting. The dominant nature of the GoF variants is well described for related Nav and Cav channels (Ortner et al. 2023; Michiels et al. 2005), but it is nonetheless intriguing that NALCN-associated disease-causing missense variants almost exclusively give rise to GoF (Monteil et al. 2023; Kschonsak et al. 2020) (Fig. 2 and 4). This is in contrast to a recent unbiased mutational screen on the related Nav1.2 channel, in which the vast majority of variants resulted in a LoF phenotype (Pablo et al. 2023). Additionally, it is worth noting that the stability distribution of the NALCN GoF variants (Fig. 4B) is broader than usual, although in line with recent work highlighting the challenges of computational models including stability predictors in identifying GoF variants (Stein et al. 2023). Together, this might indicate that NALCN has a more general propensity for variants to result in GoF phenotypes, particularly for variants located around the pore domain of NALCN. In line with previous findings, the GoF variants studied here all cluster around the lower S5 / S6, as well as some of the membrane-proximal linkers (Fig. 2A) (Kschonsak et al. 2020). Intriguingly, such spatial clustering has more generally been observed for pathogenic GoF variants compared with LoF variants in other proteins (Gerasimavicius et al. 2022; Gerasimavicius and Marsh 2025). In the case of NALCN we propose that these pore-proximal variants interfere with the stability of the channel closed state and thus result in extended open times (Kschonsak et al. 2020).

Perhaps the most intriguing finding among the GoF variants is that the functional phenotype of K1115N is only apparent when assessing the +/-Ca^2+^/Mg^2+^ ratio, while *I*_max_, AUC, and *I*_ss_ / *I*_ins_ ratio are WT-like. Given the location of K1115 at the selectivity filter (Kschonsak et al. 2020) and its previously established role in direct block by divalent cations (Chua et al. 2020), this suggests that the K1115N variant is pathogenic because of a disinhibition by divalent cations, mainly Ca^2+^ and Mg^2+^. The weaker block by these ions, especially Ca^2+^, would arguably also result in increased inward leak currents in a cellular setting, thereby leading to excessive membrane depolarization in line with a NALCN GoF. Although it is known that mutations at and around the selectivity filter of related Nav and Cav channels can cause disease, this is typically assumed to be due to a loss of selectivity or decreased conductance (Akbari Ahangar et al. 2024; Petrovicova et al. 2017; Denier et al. 2001; Geerlings et al. 2011; Bürk et al. 2014; Mangano et al. 2022; Echevarria-Cooper et al. 2022). The disinhibition by divalent cations that we propose to underlie the pathogenic nature of K1115N therefore likely represents an as of yet undescribed molecular mechanism for pathogenicity in this channel superfamily.

### Outlook

Although disease-causing variants in proteins constituting the NALCN channelosome are ultra-rare, recent years have seen a steady increase in diagnosed cases (www.channelinghope.com). Future studies will determine if the IHPRF2 syndrome (OMIM #616801), caused by autosomal recessive missense or nonsense variants of the *UNC80* gene (Monteil et al. 2023) and a less studied and yet unclassified group of patient variants presumably caused by variants in UNC79 (Bayat et al. 2023) can be subjected to the same experimental and computational framework to assess variant pathogenicity.

## Conclusions

In this study, we combined functional and computational methods to establish an ACMG-compatible framework for characterization of NALCN patient variants. Using commonly occurring NALCN variants and previously characterized patient variants, we demonstrated that our assay is able to confidently distinguish between WT-like, LoF, and GoF phenotypes. Characterizing previously untested patient variants, we found that CLIFAHDD-associated variants exlusive lead to GoF, whereas IHPRF1-associated variants lead to either LoF (for nonsense or deletion variants) or WT-like phenotypes (for missense variants). While the GoF of most CLIFAHDD variants was apparent as generally increased currents, K1115N presented a so far unique GoF mechanism based on reduced inhibition by extracellular Ca^2+^ and Mg^2+^, likely due to its position in the selectivity filter of the channel. Furthermore, protein stability predictions were able to broadly cluster NALCN variants in agreement with their functional phenotypes, providing a complementary and orthogonal tool for variant characterization.

Overall, we provide detailed insights into the molecular nature of NALCN variants that relies on two orthogonal assays, an approach we suggest be adopted to lend additional confidence to future studies into the mechanisms of NALCN-linked disease variants.

## Declarations

### Availability of data and materials

All data will be deposited in a public repository prior to publication.

## Author contributions

Acquisition of functional data: CW, HH, AA, TG; Analysis of functional data: HH, HCC; Curation and analysis of computational data: AP, AS; Writing and editing: HH, AP, AS, HCC, SAP; Overall study design and funding: AS, HCC, SAP.

## References

Abildgaard, Amanda B, Amelie Stein, Sofie V Nielsen, et al. 2019. ‘Computational and Cellular Studies Reveal Structural Destabilization and Degradation of MLH1 Variants in Lynch Syndrome’. eLife 8 (November): e49138. 10.7554/eLife.49138.

Akbari Ahangar, Amin, Eslam Elhanafy, Hayden Blanton, and Jing Li. 2024. ‘Mapping Structural Distribution and Gating-Property Impacts of Disease-Associated Mutations in Voltage-Gated Sodium Channels’. iScience 27 (9): 110678. 10.1016/j.isci.2024.110678.

Al-Sayed, Moeenaldeen D., Hamad Al-Zaidan, AlBandary Albakheet, et al. 2013. ‘Mutations in NALCN Cause an Autosomal-Recessive Syndrome with Severe Hypotonia, Speech Impairment, and Cognitive Delay’. The American Journal of Human Genetics 93 (4): 4. 10.1016/j.ajhg.2013.08.001.

Aoyagi, Kyota, Elsa Rossignol, Fadi F. Hamdan, et al. 2015. ‘A Gain-of-Function Mutation in NALCN in a Child with Intellectual Disability, Ataxia, and Arthrogryposis’. Human Mutation 36 (8): 8. 10.1002/humu.22797.

Bayat, Allan, Zhenjiang Liu, Sheng Luo, et al. 2023. ‘A New Neurodevelopmental Disorder Linked to Heterozygous Variants in UNC79’. Genetics in Medicine, May 11, 100894. 10.1016/j.gim.2023.100894.

Bend, Eric G., Yue Si, David A. Stevenson, et al. 2016. ‘NALCN Channelopathies: Distinguishing Gain-of-Function and Loss-of-Function Mutations’. Article. Neurology 87 (11): 11. 10.1212/WNL.0000000000003095.

Borovikov, A. O., I. V. Sharkova, O. P. Ryzhkova, et al. 2019. ‘Clinical and Genetic Characteristics of the Syndrome of Contractures of the Limbs and Face, Hypothony and Psychomotor Retardation (OMIM: 616 266), Caused by Mutations in the NALCN Gene’. Neuromuscular Diseases 9 (1): 83–91. 10.17650/2222-8721-2019-9-1-83-91.

Bouasse, Malik, Hathaichanok Impheng, Zoe Servant, Philippe Lory, and Arnaud Monteil. 2019. ‘Functional Expression of CLIFAHDD and IHPRF Pathogenic Variants of the NALCN Channel in Neuronal Cells Reveals Both Gain- and Loss-of-Function Properties’. Scientific Reports 9 (1): 1. 10.1038/s41598-019-48071-x.

Bramswig, Nuria C., Aida M. Bertoli-Avella, Beate Albrecht, et al. 2018. ‘Genetic Variants in Components of the NALCN–UNC80–UNC79 Ion Channel Complex Cause a Broad Clinical Phenotype (NALCN Channelopathies)’. Human Genetics 137 (9): 753–68. 10.1007/s00439-018-1929-5.

Brnich, Sarah E., Ahmad N. Abou Tayoun, Fergus J. Couch, et al. 2019. ‘Recommendations for Application of the Functional Evidence PS3/BS3 Criterion Using the ACMG/AMP Sequence Variant Interpretation Framework’. Genome Medicine 12 (1): 3. 10.1186/s13073-019-0690-2.

Bürk, Katrin, Frank J. Kaiser, Stephanie Tennstedt, et al. 2014. ‘A Novel Missense Mutation in CACNA1A Evaluated by in Silico Protein Modeling Is Associated with Non-Episodic Spinocerebellar Ataxia with Slow Progression’. European Journal of Medical Genetics 57 (5): 207–11. 10.1016/j.ejmg.2014.01.005.

Cagiada, Matteo, Kristoffer E Johansson, Audrone Valanciute, et al. 2021. ‘Understanding the Origins of Loss of Protein Function by Analyzing the Effects of Thousands of Variants on Activity and Abundance’. Molecular Biology and Evolution 38 (8): 3235–46. 10.1093/molbev/msab095.

Cagiada, Matteo, Nicolas Jonsson, and Kresten Lindorff-Larsen. 2025. ‘Decoding Molecular Mechanisms for Loss-of-Function Variants in the Human Proteome’. Preprint, bioRxiv, June 22. 10.1101/2024.05.21.595203.

Campbell, Jamie, David R. FitzPatrick, Tara Azam, et al. 2018. ‘NALCN Dysfunction as a Cause of Disordered Respiratory Rhythm With Central Apnea’. Pediatrics 141 (Supplement_5): S485–90. 10.1542/peds.2017-0026.

Chen, Ethan A., and Lauren L. Porter. 2023. ‘SSDraw: Software for Generating Comparative Protein Secondary Structure Diagrams’. Protein Science 32 (12): e4836. 10.1002/pro.4836.

Chong, Jessica X., Margaret J. McMillin, Kathryn M. Shively, et al. 2015. ‘De Novo Mutations in NALCN Cause a Syndrome Characterized by Congenital Contractures of the Limbs and Face, Hypotonia, and Developmental Delay’. The American Journal of Human Genetics 96 (3): 3. 10.1016/j.ajhg.2015.01.003.

Chua, H. C., M. Wulf, C. Weidling, L. P. Rasmussen, and S. A. Pless. 2020. ‘The NALCN Channel Complex Is Voltage Sensitive and Directly Modulated by Extracellular Calcium’. Research Article. Science Advances 6 (17): 17. 10.1126/sciadv.aaz3154.

Denier, Christian, Anne Ducros, Alexandra Durr, Bruno Eymard, Bénédicte Chassande, and Elisabeth Tournier-Lasserve. 2001. ‘Missense CACNA1A Mutation Causing Episodic Ataxia Type 2’. Archives of Neurology 58 (2): 292–95. 10.1001/archneur.58.2.292.

Echevarria-Cooper, Dennis M, Nicole A Hawkins, Sunita N Misra, et al. 2022. ‘Cellular and Behavioral Effects of Altered NaV1.2 Sodium Channel Ion Permeability in Scn2aK1422E Mice’. Human Molecular Genetics 31 (17): 2964–88. 10.1093/hmg/ddac087.

Frenz, Brandon, Steven M. Lewis, Indigo King, Frank DiMaio, Hahnbeom Park, and Yifan Song. 2020. ‘Prediction of Protein Mutational Free Energy: Benchmark and Sampling Improvements Increase Classification Accuracy’. Frontiers in Bioengineering and Biotechnology 8 (October). 10.3389/fbioe.2020.558247.

Geerlings, Rianne PJ, Peter J Koehler, Danielle YP Haane, et al. 2011. ‘Head Tremor Related to CACNA1A Mutations’. Cephalalgia 31 (12): 1315–19. 10.1177/0333102411414442.

Gerasimavicius, Lukas, Benjamin J. Livesey, and Joseph A. Marsh. 2022. ‘Loss-of-Function, Gain-of-Function and Dominant-Negative Mutations Have Profoundly Different Effects on Protein Structure’. Nature Communications 13 (1): 3895. 10.1038/s41467-022-31686-6.

Gerasimavicius, Lukas, and Joseph A. Marsh. 2025. ‘A Knowledge-Based Distance Metric Highlights Underperformance of Variant Effect Predictors on Gain-of-Function Missense Variants’. Preprint, bioRxiv, July 28. 10.1101/2025.07.23.666325.

Hadouiri, Nawale, Lidiane Pereira Garcia, Romain Baudat, et al. 2025. ‘De Novo Variants of NALCN Differentially Impact Both the Phenotypic Spectrum of Patients and the Biophysical Properties of the NALCN Current’. Preprint, medRxiv, June 22. 10.1101/2025.06.20.25329825.

Høie, Magnus Haraldson, Matteo Cagiada, Anders Haagen Beck Frederiksen, Amelie Stein, and Kresten Lindorff-Larsen. 2022. ‘Predicting and Interpreting Large-Scale Mutagenesis Data Using Analyses of Protein Stability and Conservation’. Cell Reports 38 (2). 10.1016/j.celrep.2021.110207.

Jumper, John, Richard Evans, Alexander Pritzel, et al. 2021. ‘Highly Accurate Protein Structure Prediction with AlphaFold’. Nature 596 (7873): 583–89. 10.1038/s41586-021-03819-2.

Kang, Yunlu, and Lei Chen. 2022. ‘Structure and Mechanism of NALCN-FAM155A-UNC79-UNC80 Channel Complex’. Nature Communications 13 (1): 1. 10.1038/s41467-022-30403-7.

Kang, Yunlu, Jing-Xiang Wu, and Lei Chen. 2020. ‘Structure of Voltage-Modulated Sodium-Selective NALCN-FAM155A Channel Complex’. Nature Communications 11 (1): 1. 10.1038/s41467-020-20002-9.

Karczewski, Konrad J., Laurent C. Francioli, Grace Tiao, et al. 2020. ‘The Mutational Constraint Spectrum Quantified from Variation in 141,456 Humans’. Nature 581 (7809): 434–43. 10.1038/s41586-020-2308-7.

Köroğlu, Çiğdem, Mehmet Seven, and Aslihan Tolun. 2013. ‘Recessive Truncating NALCN Mutation in Infantile Neuroaxonal Dystrophy with Facial Dysmorphism’. Journal of Medical Genetics 50 (8): 8. 10.1136/jmedgenet-2013-101634.

Kschonsak, Marc, Han Chow Chua, Cameron L. Noland, et al. 2020. ‘Structure of the Human Sodium Leak Channel NALCN’. Nature, July 22, 1–10. 10.1038/s41586-020-2570-8.

Kschonsak, Marc, Han Chow Chua, Claudia Weidling, et al. 2022. ‘Structural Architecture of the Human NALCN Channelosome’. Nature 603 (7899): 180–86. 10.1038/s41586-021-04313-5.

Larsen-Ledet, Sven, Aleksandra Panfilova, and Amelie Stein. 2025. ‘Disentangling the Mutational Effects on Protein Stability and Interaction of Human MLH1’. PLOS Genetics 21 (4): e1011681. 10.1371/journal.pgen.1011681.

Lee, Jung-Ha, Leanne L. Cribbs, and Edward Perez-Reyes. 1999. ‘Cloning of a Novel Four Repeat Protein Related to Voltage-Gated Sodium and Calcium Channels’. FEBS Letters 445 (2–3): 2–3. 10.1016/S0014-5793(99)00082-4.

Lu, Boxun, Yanhua Su, Sudipto Das, Jin Liu, Jingsheng Xia, and Dejian Ren. 2007. ‘The Neuronal Channel NALCN Contributes Resting Sodium Permeability and Is Required for Normal Respiratory Rhythm’. Cell 129 (2): 2. 10.1016/j.cell.2007.02.041.

Ma, Joanne G., Matthew J. O’Neill, Ebony Richardson, et al. 2024. ‘Multisite Validation of a Functional Assay to Adjudicate SCN5A Brugada Syndrome–Associated Variants’. Circulation: Genomic and Precision Medicine 17 (4): e004569. 10.1161/CIRCGEN.124.004569.

Mangano, Giuseppe Donato, Antonina Fontana, Vincenzo Antona, et al. 2022. ‘Commonalities and Distinctions between Two Neurodevelopmental Disorder Subtypes Associated with SCN2A and SCN8A Variants and Literature Review’. Molecular Genetics & Genomic Medicine 10 (5): e1911. 10.1002/mgg3.1911.

Michiels, Jan J., Rene H. M. te Morsche, Jan B. M. J. Jansen, and Joost P. H. Drenth. 2005. ‘Autosomal Dominant Erythermalgia Associated With a Novel Mutation in the Voltage-Gated Sodium Channel **α** Subunit Nav1.7’. Archives of Neurology 62 (10): 1587–90. 10.1001/archneur.62.10.1587.

Monteil, Arnaud, Adriano Senatore, Antonio Gil-Nagel, Paloma Parra-Diaz, Philippe Lory, and Nathalie C. Guérineau. 2023. ‘New Insights into the Physiology and Pathophysiology of the Atypical Sodium Leak Channel NALCN’. Physiological Reviews, ahead of print, August 24. 10.1152/physrev.00014.2022.

Nguyen, Emmanuelle, Martine Tétreault, Dènahin Hinnoutondji Toffa, Patrick Cossette, Éric Samarut, and Dang Khoa Nguyen. 2023. ‘Novel NALCN Variant Linked to Temporal Lobe Epilepsy’. American Journal of Medical Genetics. Part A 191 (7): 1942–47. 10.1002/ajmg.a.63209.

Ortner, Nadine J., Anupam Sah, Enrica Paradiso, et al. 2023. ‘The Human Channel Gating– Modifying A749G CACNA1D (Cav1.3) Variant Induces a Neurodevelopmental Syndrome– like Phenotype in Mice’. JCI Insight 8 (20). 10.1172/jci.insight.162100.

Pablo, Juan Lorenzo B., Savannah L. Cornett, Lei A. Wang, et al. 2023. ‘Scanning Mutagenesis of the Voltage-Gated Sodium Channel NaV1.2 Using Base Editing’. Cell Reports 42 (6). 10.1016/j.celrep.2023.112563.

Park, Hahnbeom, Philip Bradley, Per Jr. Greisen, et al. 2016. ‘Simultaneous Optimization of Biomolecular Energy Functions on Features from Small Molecules and Macromolecules’. Journal of Chemical Theory and Computation 12 (12): 6201–12. 10.1021/acs.jctc.6b00819.

Parra-Díaz, Paloma, Arnaud Monteil, Daniel Calame, et al. 2025. ‘Genotype-Phenotype Landscape of NALCN and UNC80-Related Disorders’. Neurology 104 (7): e213429. 10.1212/WNL.0000000000213429.

Petrovicova, Andrea, Miroslav Brozman, Egon Kurca, et al. 2017. ‘Novel Missense Variant of CACNA1A Gene in a Slovak Family with Episodic Ataxia Type 2’. Biomedical Papers 161 (1): 107–10. 10.5507/bp.2016.066.

Rives, Alexander, Joshua Meier, Tom Sercu, et al. 2021. ‘Biological Structure and Function Emerge from Scaling Unsupervised Learning to 250 Million Protein Sequences’. Proceedings of the National Academy of Sciences 118 (15): e2016239118. 10.1073/pnas.2016239118.

Seven, M., A. Ozkiliç, and A. Yüksel. 2002. ‘Dysmorphic Face in Two Siblings with Infantile Neuroaxonal Dystrophy’. Genetic Counseling (Geneva, Switzerland) 13 (4): 465–73.

Stein, David, Meltem Ece Kars, Yiming Wu, et al. 2023. ‘Genome-Wide Prediction of Pathogenic Gain- and Loss-of-Function Variants from Ensemble Learning of a Diverse Feature Set’. Genome Medicine 15 (1): 103. 10.1186/s13073-023-01261-9.

Tehrani Fateh, Sahand, Saman Bagheri, Hossein Sadeghi, et al. 2023. ‘Extending and Outlining the Genotypic and Phenotypic Spectrum of Novel Mutations of NALCN Gene in IHPRF1 Syndrome: Identifying Recurrent Urinary Tract Infection’. Neurological Sciences: Official Journal of the Italian Neurological Society and of the Italian Society of Clinical Neurophysiology 44 (12): 4491–98. 10.1007/s10072-023-06960-0.

Thompson, Christopher H., Franck Potet, Tatiana V. Abramova, et al. 2023. ‘Epilepsy-Associated SCN2A (NaV1.2) Variants Exhibit Diverse and Complex Functional Properties’. Journal of General Physiology 155 (10): e202313375. 10.1085/jgp.202313375.

Tiemann, Johanna Katarina Sofie, Henrike Zschach, Kresten Lindorff-Larsen, and Amelie Stein. 2023. ‘Interpreting the Molecular Mechanisms of Disease Variants in Human Transmembrane Proteins’. Biophysical Journal 122 (11): 2176–91. 10.1016/j.bpj.2022.12.031.

Vanoye, Carlos G., Reshma R. Desai, Zhigang Ji, et al. 2022. ‘High-Throughput Evaluation of Epilepsy-Associated KCNQ2 Variants Reveals Functional and Pharmacological Heterogeneity’. JCI Insight 7 (5). 10.1172/jci.insight.156314.

Vivero, M., M.t. Cho, A. Begtrup, et al. 2017. ‘Additional de Novo Missense Genetic Variants in NALCN Associated with CLIFAHDD Syndrome’. Clinical Genetics 91 (6): 929–31. 10.1111/cge.12899.

Xie, Jiongfang, Meng Ke, Lizhen Xu, et al. 2020. ‘Structure of the Human Sodium Leak Channel NALCN in Complex with FAM155A’. Nature Communications 11 (1): 1. 10.1038/s41467-020-19667-z.

Zhou, Lunni, Haobin Liu, Qingqing Zhao, Jianping Wu, and Zhen Yan. 2022. ‘Architecture of the Human NALCN Channelosome’. Cell Discovery 8 (1): 33. 10.1038/s41421-022-00392-4.

