## Supplementary Information for "Framework for combined functional and computational assessment of variant pathogenicity in the sodium leak channel NALCN"

*Table S1: Data overview for electrophysiological recordings shown in figures 2 and 3 with Imax, AUC, $I_{ss}/I_{ins}$ Ratio and +/-$Ca^{2+}$/$Mg^{2+}$ Ratio. Values are given as mean ± SD (n). n/a: not applicable.*

| Figure 2 | | | | |
|---|---|---|---|---|
| Construct | $I_{max}$ (/WT) | AUC (/WT) | $I_{ss}/I_{ins}$ Ratio | +/-$Ca^{2+}$/$Mg^{2+}$ Ratio |
| WT | 1.00 ± 0.19 (78) | 1.00 ± 0.26 (69) | 0.18 ± 0.03 (92) | 0.18 ± 0.05 (92) |
| V1020F | 3.10 ± 0.97 (11) | 11.84 ± 3.48 (11) | 0.86 ± 0.04 (11) | 0.17 ± 0.02 (11) |
| W1287L | 0.24 ± 0.08 (8) | 0.22 ± 0.10 (8) | n/a | n/a |
| V70L | 1.10 ± 0.25 (8) | 0.96 ± 0.22 (8) | 0.18 ± 0.02 (8) | 0.16 ± 0.01 (8) |
| R372H | 1.11 ± 0.09 (8) | 0.90 ± 0.12 (8) | 0.17 ± 0.02 (8) | 0.16 ± 0.02 (8) |
| H769Y | 1.03 ± 0.26 (18) | 1.08 ± 0.24 (10) | 0.16 ± 0.03 (10) | 0.21 ± 0.06 (10) |
| A1091E | 1.18 ± 0.20 (8) | 1.01 ± 0.06 (8) | 0.16 ± 0.01 (8) | 0.16 ± 0.02 (8) |
| K1115N hom | 0.85 ± 0.22 (12) | 1.12 ± 0.28 (12) | 0.26 ± 0.05 (12) | 0.53 ± 0.05 (12) |
| K1115N het | 0.95 ± 0.17 (12) | 1.04 ± 0.20 (12) | 0.20 ± 0.02 (12) | 0.43 ± 0.02 (12) |
| F1141V hom | 2.94 ± 1.81 (17) | 4.46 ± 3.72 (19) | 0.23 ± 0.05 (19) | 0.15 ± 0.02 (19) |
| F1141V het | 2.17 ± 0.90 (19) | 3.16 ± 1.57 (19) | 0.20 ± 0.03 (19) | 0.16 ± 0.02 (19) |
| L1150V hom | 2.75 ± 0.87 (26) | 4.31 ± 1.41 (26) | 0.33 ± 0.11 (21) | 0.18 ± 0.05 (26) |
| L1150V het | 2.32 ± 0.62 (17) | 3.44 ± 1.61 (17) | 0.33 ± 0.10 (12) | 0.18 ± 0.04 (17) |
| R1174G hom | 3.31 ± 1.04 (15) | 9.21 ± 3.58 (15) | 0.60 ± 0.08 (11) | 0.25 ± 0.09 (15) |
| R1174G het | 3.15 ± 0.64 (13) | 7.92 ± 2.56 (13) | 0.59 ± 0.16 (13) | 0.23 ± 0.10 (13) |
| V1328M hom | 3.51 ± 1.90 (19) | 6.23 ± 3.83 (19) | 0.29 ± 0.05 (19) | 0.16 ± 0.02 (19) |
| V1328M het | 2.67 ± 2.14 (16) | 5.49 ± 6.10 (16) | 0.29 ± 0.08 (14) | 0.16 ± 0.04 (16) |
| I1445L hom | 3.75 ± 1.03 (14) | 7.38 ± 3.26 (15) | 0.39 ± 0.21 (14) | 0.23 ± 0.09 (15) |
| I1445L het | 1.89 ± 0.54 (14) | 2.76 ± 1.10 (14) | 0.25 ± 0.07 (14) | 0.20 ± 0.03 (14) |
| L1452S hom | 2.78 ± 0.86 (16) | 5.05 ± 2.45 (16) | 0.50 ± 0.13 (11) | 0.19 ± 0.05 (16) |
| L1452S het | 2.66 ± 0.60 (13) | 5.09 ± 2.55 (13) | 0.51 ± 0.18 (8) | 0.22 ± 0.07 (13) |
| Figure 3 | | | | |
| Construct | $I_{max}$ (/WT) | AUC (/WT) | $I_{ss}/I_{ins}$ Ratio | +/-$Ca^{2+}$/$Mg^{2+}$ Ratio |
| WT | 1.02 ± 0.28 (43) | 1.00 ± 0.26 (43) | n/a | n/a |
| W107* | 0.44 ± 0.04 (3) | 0.55 ± 0.05 (3) | n/a | n/a |
| W179* | 0.40 ± 0.11 (9) | 0.58 ± 0.16 (9) | n/a | n/a |
| Q642* | 0.30 ± 0.06 (7) | 0.43 ± 0.15 (7) | n/a | n/a |
| R1008* | 0.42 ± 0.09 (7) | 0.65 ± 0.20 (7) | n/a | n/a |
| Q1186* | 0.40 ± 0.09 (9) | 0.58 ± 0.16 (9) | n/a | n/a |
| R1384* | 0.34 ± 0.09 (7) | 0.44 ± 0.11 (7) | n/a | n/a |
| V891Sfs*2 | 0.29 ± 0.11 (6) | 0.48 ± 0.11 (6) | n/a | n/a |
| ΔV956-L963 | 0.30 ± 0.10 (6) | 0.46 ± 0.08 (6) | n/a | n/a |
| ΔV1020-R1054 | 0.52 ± 0.16 (8) | 0.59 ± 0.25 (8) | n/a | n/a |
| G883V | 1.26 ± 0.33 (8) | 1.22 ± 0.34 (8) | n/a | n/a |
| WT | 1.00 ± 0.22 (14) | 1.00 ± 0.33 (14) | n/a | n/a |
| W37* | 0.06 ± 0.00 (5) | 0.13 ± 0.04 (5) | n/a | n/a |
| P908L | 0.72 ± 0.22 (10) | 0.70 ±0.38 (10) | n/a | n/a |
| W37*/P908L | 0.74 ± 0.33 (7) | 0.67 ± 0.34 (7) | n/a | n/a |
| WT | 1.00 ± 0.24 (11) | 1.00 ± 0.32 (16) | n/a | n/a |
| R855* | 0.06 ± 0.01 (3) | 0.15 ± 0.02 (3) | n/a | n/a |
| R1094Q | 0.70 ± 0.23 (9) | 0.57 ± 0.23 (10) | n/a | n/a |
| R855*/R1094Q | 0.66 ± 0.48 (7) | 1.03 ± 0.99 (7) | n/a | n/a |

*Figure S1: Scatter plots of the change in protein stability (ΔΔG in kcal/mol, x axis) and the estimated sequence tolerance (arbitrary units, y axis) for individual positions highlighted in Figure 4B. Colored data points denote amino acid exchanges that have been reported as common variants (green) or patient variants (pink for previously tested CLIFAHDD variants, red for CLIFAHDD variants tested in this study; blue for homozygous IHPRF1 variants, purple for compound heterozygous IHPRF1 variants), whereas grey data points represent all other possible exchanges for that position. By convention, positive ΔΔGs denote destabilising variants, while negative ΔΔGs indicate stabilising variants.*

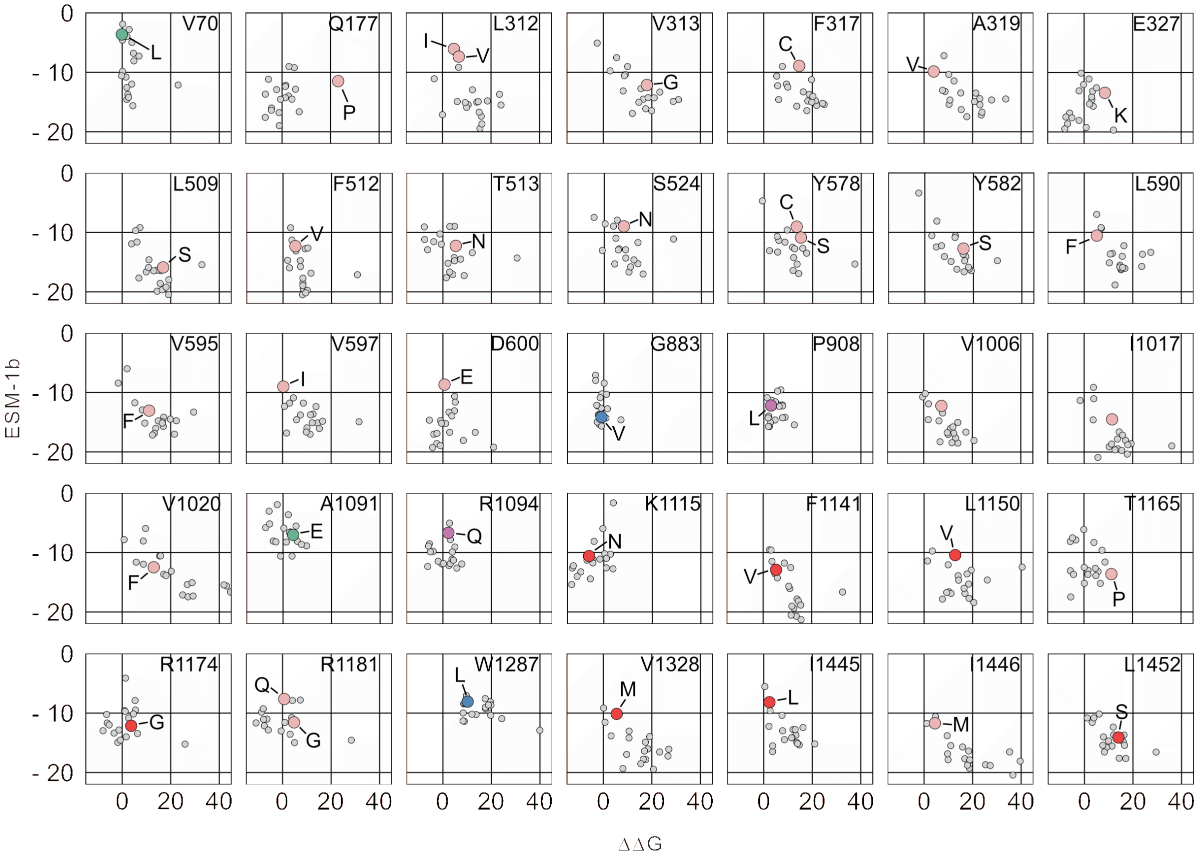
